# Identification of 14 Known Drugs as Inhibitors of the Main Protease of SARS-CoV-2

**DOI:** 10.1101/2020.08.28.271957

**Authors:** Mohammad M. Ghahremanpour, Julian Tirado-Rives, Maya Deshmukh, Joseph A. Ippolito, Chun-Hui Zhang, Israel Cabeza de Vaca, Maria-Elena Liosi, Karen S. Anderson, William L. Jorgensen

**Affiliations:** Department of Chemistry, Yale University, New Haven, Connecticut 06520-8107; Department of Pharmacology, Yale University School of Medicine, New Haven, CT 06520-8066; Department of Molecular Biophysics and Biochemistry, Yale University School of Medicine, New Haven, CT 06520-8066

## Abstract

A consensus virtual screening protocol has been applied to ca. 2000 approved drugs to seek inhibitors of the main protease (M^pro^) of SARS-CoV-2, the virus responsible for COVID-19. 42 drugs emerged as top candidates, and after visual analyses of the predicted structures of their complexes with M^pro^, 17 were chosen for evaluation in a kinetic assay for M^pro^ inhibition. Remarkably 14 of the compounds at 100-μM concentration were found to reduce the enzymatic activity and 5 provided IC_50_ values below 40 μM: manidipine (4.8 μM), boceprevir (5.4 μM), lercanidipine (16.2 μM), bedaquiline (18.7 μM), and efonidipine (38.5 μM). Structural analyses reveal a common cloverleaf pattern for the binding of the active compounds to the P1, P1’, and P2 pockets of M^pro^. Further study of the most active compounds in the context of COVID-19 therapy is warranted, while all of the active compounds may provide a foundation for lead optimization to deliver valuable chemotherapeutics to combat the pandemic.

## INTRODUCTION

SARS-CoV-2, the cause of the COVID-19 pandemic,^1^ is a coronavirus (CoV) from the *Coronaviridae* family. Its RNA genome is ~82% identical to that of SARS-CoV,^2^ which was responsible for the Severe Acute Respiratory Syndrome (SARS) pandemic in 2003.^3^ SARS-CoV-2 encodes two cysteine proteases: the chymotrypsin-like cysteine or main protease, known as 3CL^pro^ or M^pro^, and the papain-like cysteine protease, PL^pro^. They catalyze the proteolysis of polyproteins translated from the viral genome into non-structural proteins essential for packaging the nascent virion and viral repication.^4^ Therefore, inhibiting the activity of these proteases would impede the replication of the virus. M^pro^ processes the polyprotein 1ab at multiple cleavage sites. It hydrolyzes the Gln-Ser peptide bond in the Leu-Gln-Ser-Ala-Gly recognition sequence. This cleavage site in the substrate is distinct from the peptide sequence recognized by other human cysteine proteases known to date.^5^ Thus, M^pro^ is viewed as a promising target for anti SARS-CoV-2 drug design; it has been the focus of several studies since the pandemic has emerged. ^2,4–7^

An X-ray crystal structure of M^pro^ reveals that it forms a homodimer with a 2-fold crystallographic symmetry axis.^2,5^ Each protomer, with a length of 306 residues, is made of three domains (I-III). Domains II and I fold into a six-stranded β-barrel that harbors the active site.^2,4,5^ Domain III forms a cluster of five antiparallel α-helices that regulates the dimerization of the protease. A flexible loop connects domain II to domain III. The M^pro^ active site contains a Cys-His catalytic dyad and canonical binding pockets that are denoted P1, P1′, P2, P3, and P4.^2^ The amino acid sequence of the active site is highly conserved among coronaviruses.^8^ The catalytic dyad residues are His^41^ and Cys^145^ and the residues playing key roles in the binding of the substrate are Phe^140^, His^163^, Met^165^, Glu^166^, and Gln^189^ (Figure 1). These residues have been found to interact with the ligands co-crystallized with M^pro^ in different studies.^2,4,5^ Crystallographic data also suggested that Ser^1^ of one protomer interacts with Phe^140^ and Glu^166^ of the other as the result of dimerization.^2,4^ These interactions stabilize the P1 binding pocket, thereby, dimerization of the main protease is likely for its catalytic activity.^2,4^

**Figure 1.**
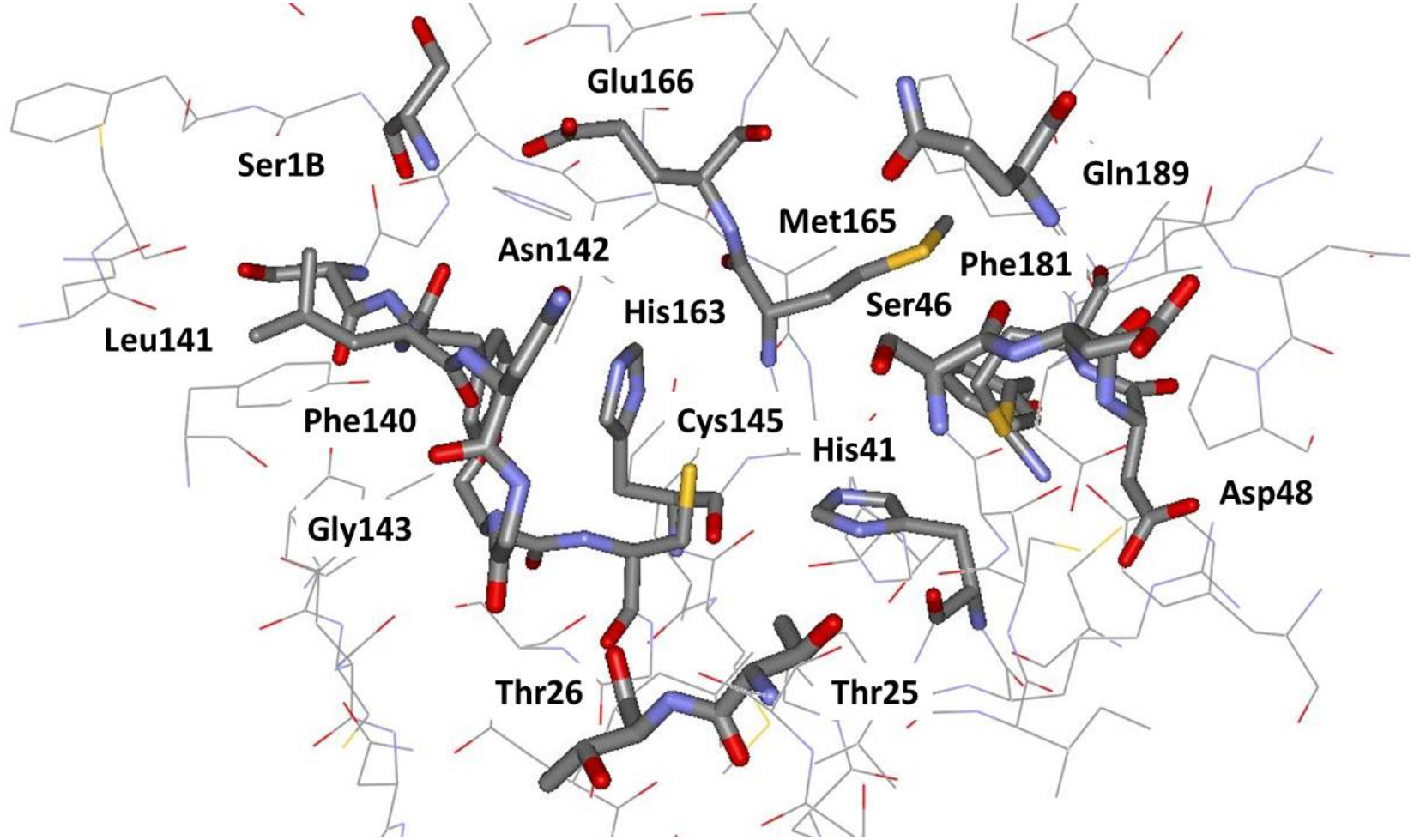
Rendering of the residues near the catalytic site of M^Pro^ from a crystal structure at 1.31-Å resolution (PDB ID: 5R82). The catalytic residues are His41 and Cys145.

Drug repurposing is an important strategy for immediate response to the COVID-19 pandemic.^9^ In this approach, the main goal of computational and experimental studies has been to find existing drugs that might be effective against SARS-CoV-2. For instance, a molecular docking study suggested remdesivir as a potential therapeutic that could be used against SARS-CoV-2,^10^ which was supported experimentally by an EC_50_ value of 23 μM in an infected-cell assay.^11^ However, a clinical trial showed no statistically significant clinical benefits of remdesivir on adult patients hospitalized for severe COVID-19.^12^ Nonetheless, patients who were administered remdesivir in the same trial showed a faster time to clinical improvement in comparison to the placebo-control group.^12^ In another clinical trial, only patients on mechanical ventilation benefitted from remdesivir.^13^ An EC_50_ value of 27 μM was also reported for lopinavir^11^, suggesting it may have beneficial activity against SARS-CoV-2. However, neither lopinavir nor the lopinavir/ritonavir combination has thus far shown any significant benefits against COVID-19 in clinical trials. Chloroquine, hydroxychloroquine, and favipiravir have also been explored for repurposing against COVID-19; however, clinical studies with them have been controversial.^14–17^ These studies reflect the urgent need for systematic drug discovery efforts for therapies effective against SARS-CoV-2.

Thus, we decided to pursue discovery of small-molecule inhibitors of M^pro^. The aim of this initial work was two-fold: to identify known drugs that may show some activity, but also to identify structurally promising, synthetically-accessible substructures suitable for subsequent lead optimization. Our expectation was that existing drugs may show activity but not at the low-nanomolar levels that are typical of effective therapies. This report provides results for the first goal. The work began by designing and executing a consensus molecular docking protocol to virtual screen ~2000 approved drugs. The predicted structures (poses) of the complexes for the top-scoring 42 drugs received extensive scrutiny including consideration of intermolecular contacts, conformation, stability in molecular dynamics (MD) simulations, and potential for synthetic modification to arrive at 17 drugs, which were purchased and assayed for inhibition of M^pro^. The outcome was strikingly successful with 14 of the 17 compounds showing some reduction of M^pro^ activity at 100 μM concentration, and with 5 compounds yielding IC_50_ values below 40 μM. The most potent inhibitors of M^pro^ identified here, manidipine and boceprevir, have IC_50_ values of 4.8 and 5.4 μM, respectively.

## COMPUTATIONAL APPROACH

### Selection of the Crystal Structure of M^pro^

Our analyses of more than 50 crystal structures of SARS-CoV-2 main protease in apo and holo forms showed small structural variations in the active site region. The overall root mean square deviation (RMSD) of all structures was ~0.8 Å for C_α_ atoms. The presence of a ligand in the crystal structure likely places the side chains of the active site residues in positions that are more suitable for performing molecular docking compared to the apo form of the enzyme. Thus, we chose to use a high-resolution (1.31 Å) structure of M^pro^ co-crystalized with a non-covalent small fragment hit (PDB ID 5R82)^18^ for docking the approved drugs after removal of the fragment (Figure 1). The program Reduce^19^ was run on the structure for allowing side-chain flips, optimizing hydrogen bonds, and adding/removing hydrogen atoms. The p*K*a values of the ionizable residues of M^pro^ were predicted using the PROPKA3^20^ and the H++ severs.^21,22^ Accordingly, lysines and arginines were positively charged, aspartic and glutamic acids were negatively charged, and all histidines were neutral. All histidines were built with the proton on Nε except for His80, which was protonated at Nδ. The resulting M^pro^ structure has a net charge of −4 *e*. Extensive visual inspection was carried out using UCSF Chimera.^23^

### Consensus Molecular Docking

Most docking programs apply methods to generate an initial set of conformations, and tautomeric and protonation states for each ligand. This is followed by application of search algorithms and scoring functions to generate and score the poses of the ligand in the binding site of a protein. Scoring functions have been trained to reproduce a finite set of experimental ligand-binding affinities that are generally a mix of activity data converted to a free-energy scale. Therefore, the accuracy of the scores is dependent on multiple factors including the compounds that were part of the training set. To mitigate the biases, we performed four independent runs of protein-ligand docking with a library of ca. 2000 approved, oral drugs using Glide, AutoDock Vina, and two protocols with AutoDock 4.2. The results were compiled and further consideration focused on those compounds that ranked among the top 10% percent in at least 3 out of the 4 runs.

### Glide

Schrödinger’s Protein Prep wizard utility was used for preparing the protein. A 20-Å grid was then generated and centered on the co-crystallized ligand, which was subsequently removed. The drug library members were neutralized and/or ionized via Schrödinger’s LigPrep.^24^ The Epik program^25^ was used for estimating the p*K*a values of each compound. Plausible tautomers and stereoisomers within the pH range of 7 ±1 were generated for each compound using the OPLS3 force field.^26^ These conditions resulted in a total of 16000 structures, which were then docked into M^pro^ using Schrödinger’s standard-precision (SP) Glide.^27,28^

### AutoDock

The AutoDockTools (ADT) software^29^ was used for creating PDBQT files from SDF and PDB files of compounds and the protein, respectively. Non-polar hydrogen atoms were removed and Gasteiger–Marsili charges were assigned for both the protein and the ligands using ADT. The AutoGrid 4.2 program^29^ was used for generating affinity grids with a spacing of 0.375 Å and with a box size of 74 × 80 × 62 Å. The affinity grids were centered at two different points of the active site for performing two sets of runs. In the first run, the grid box was centered at C_β_ of Cys145 of monomer A. In the second run, the grid center was displaced toward the geometric center of the active site. The AutoDock 4.2 program^29^ was applied for docking the ligands into M^pro^. The Lamarckian genetic algorithm (LGA) was used for ligand conformational searching. LGA was iterated 15 and 50 times in the first and the second run, respectively, for each compound. The maximum RMS tolerance for conformational cluster analysis was 2.0 and 0.5 Å in the first and second runs, respectively. The number of generations was set to 27000 with 300 individuals in each population in both runs. The maximum number of energy evaluations was 30 × 10^6^ for all compounds and 40 × 10^6^ for re-docking of the selected consensus compounds. Other parameters were set to their default values.

### AutoDock Vina

The PDBQT files generated by ADT for the protein and library compounds also used for running AutoDock Vina.^30^ Non-polar hydrogen atoms were removed. An affinity grid box with a size of 18 × 21 × 18 Å was generated and centered on the active site. The default docking parameters were used, except for the number of modes that was set to 9.

### Molecular Dynamics Simulations

The GROMACS software, version 2018a compiled in double precision, was used for performing all molecular dynamics (MD) simulations.^31^ The protonated M^pro^ dimer, with a net charge of −8 *e*, was represented by the OPLS-AA/M force field.^32^ TIP4P water was used as the solvent.^33^ Sodium counterions were added to neutralize the net charge of each system. The selected ligand candidates were represented by the OPLS/CM1A force field,^34^ as assigned by the BOSS software^35^ (version 4.9) and the LigParGen Python code.^36^ The parameters were converted to GROMACS format using LigParGen.^36^ For neutral ligands, the CM1A partial atomic charges were scaled by a factor of 1.14.^34^

Each M^pro^-ligand complex was put at the center of a triclinic simulation box with 10-Å padding. An energy minimization was then performed until the steepest descent algorithm converged to a maximum force smaller than 2.4 kcal mol^−1^ Å^−1^. A cutoff radius of 12 Å was used to explicitly calculate non-bonded interactions. Long-range electrostatic interactions were treated using the Particle Mesh Ewald (PME) algorithm.^37^ The PME was used with an interpolation order of 4, a Fourier spacing of 1.2 Å, and a relative tolerance of 10^−6^. The van der Waals forces were smoothly switched to zero between 10 and 12 Å. Analytical corrections to the long-range effect of dispersion interactions were applied to both energy and pressure. All covalent bonds to hydrogen atoms were constrained at their equilibrium lengths using the LINCS algorithm^38^ with the order of 12 in the expansion of the constraint-coupling matrix. Each system was subsequently simulated for 1 ns in the canonical ensemble (*NVT*) in order for the solvent to relax and the temperature of the system to equilibrate. Initial velocities were sampled from a Maxwell-Boltzmann distribution at 310 K. The V-rescale thermostat with a stochastic term^39^ was used for keeping the temperature at 310 K. The stochastic term ensured that the sampled ensemble was canonical.^39^ The coupling constant of the thermostat was set to 2.0 ps. The system was then equilibrated for 1.5 ns in the isothermal-isobaric ensemble (*NPT*) for obtaining a density consistent with the reference pressure. The pressure was kept at 1 bar by the Berendsen barostat^40^ with a coupling constant of 4.0 ps and a compressibility factor of 4.5 × 10^−5^ bar. A harmonic position restraint with a force constant of 2.4 kcal mol^−1^ Å^−2^ was applied to the protein backbone and to all solute heavy atoms during the equilibration steps. A 70 ns unrestrained run was then performed in the NPT ensemble with the Parrinello-Rahman barostat^41^ using a coupling time of 4.0 ps.

## EXPERIMENTAL DETAILS

### Expression and Purification of SARS-CoV-2 M^pro^

A PGEX-6p-1 vector containing the gene for SARS-CoV-2 M^pro^ harboring a His6 tag followed by a modified PreScission cleavage site was used to produce recombinant protein.^5^ Recombinant M^pro^ with authentic N- and C-termini was expressed and purified as previously described.^5^

### Kinetic Assays of SARS-CoV-2 M^pro^ Activity and Analysis

All assayed compounds were obtained from commercial sources except cinnoxicam, which had to be synthesized, and had purity >95% based on HPLC analysis. Kinetics of SARS-CoV-2 M^pro^ were measured as previously described.^5,7^ Briefly, 100 nM M^pro^ in reaction buffer (20mM Tris, 100 mM NaCl, 1 mM DTT, pH 7.3) was incubated with or without compound in DMSO at varying concentrations to a final DMSO concentration of 6% for 15 minutes with shaking at room temperature. The reaction was initiated by addition of substrate (Dabcyl-KTSAVLQ↓SGFRKM-E(Edans-NH2); GL Biochem) in reaction buffer, which is cleaved by M^pro^, generating a product containing a free Edans group. Fluorescence was monitored at an excitation wavelength of 360 nm and emission wavelength of 460 nm. All measurements were performed in triplicate and averaged. IC_50_ plots and values were generated using Prism 8.0 (GraphPad).

## RESULTS AND DISCUSSION

### Virtual Screening

The docking scores obtained for all compounds range from −10.85 to −0.59 for Glide, from −12.33 to −2.30 for AutoDock run 1, from −10.74 to −0.40 for AutoDock run 2, and −8.50 to −2.10 for AutoDock Vina. As expected, the range of scores is wide and it is different from one docking program to another. The complete list of compounds and docking scores is provided in the SI. Compounds were ranked based on their docking scores and the top 200 hits from the four docking runs were compared. As the result, 42 compounds with a consensus count of 4 or 3 were selected. This means that these compounds were among the top-200 ranked compounds in all 4 or at least 3 out of the 4 docking runs. The indications and mechanisms of action for the 42 drugs are shown in Table 1, and the structures of some of the ones that turned out to be most interesting are shown in Figure 2. The primary indications include bacterial and viral infections, hypertension, psychosis, inflammation, and cancer. Their mechanisms of action are also broad ranging from kinase and protease inhibitors to dopamine receptors agonists/antagonists, and calcium channel blockers. It is not surprising that peptidic protease inhibitors are well-represented in view of the peptide substrate and prior discovery of peptidic inhibitors for M^pro^ and its SARS-CoV relative. ^7,42,43^

**Table 1.**
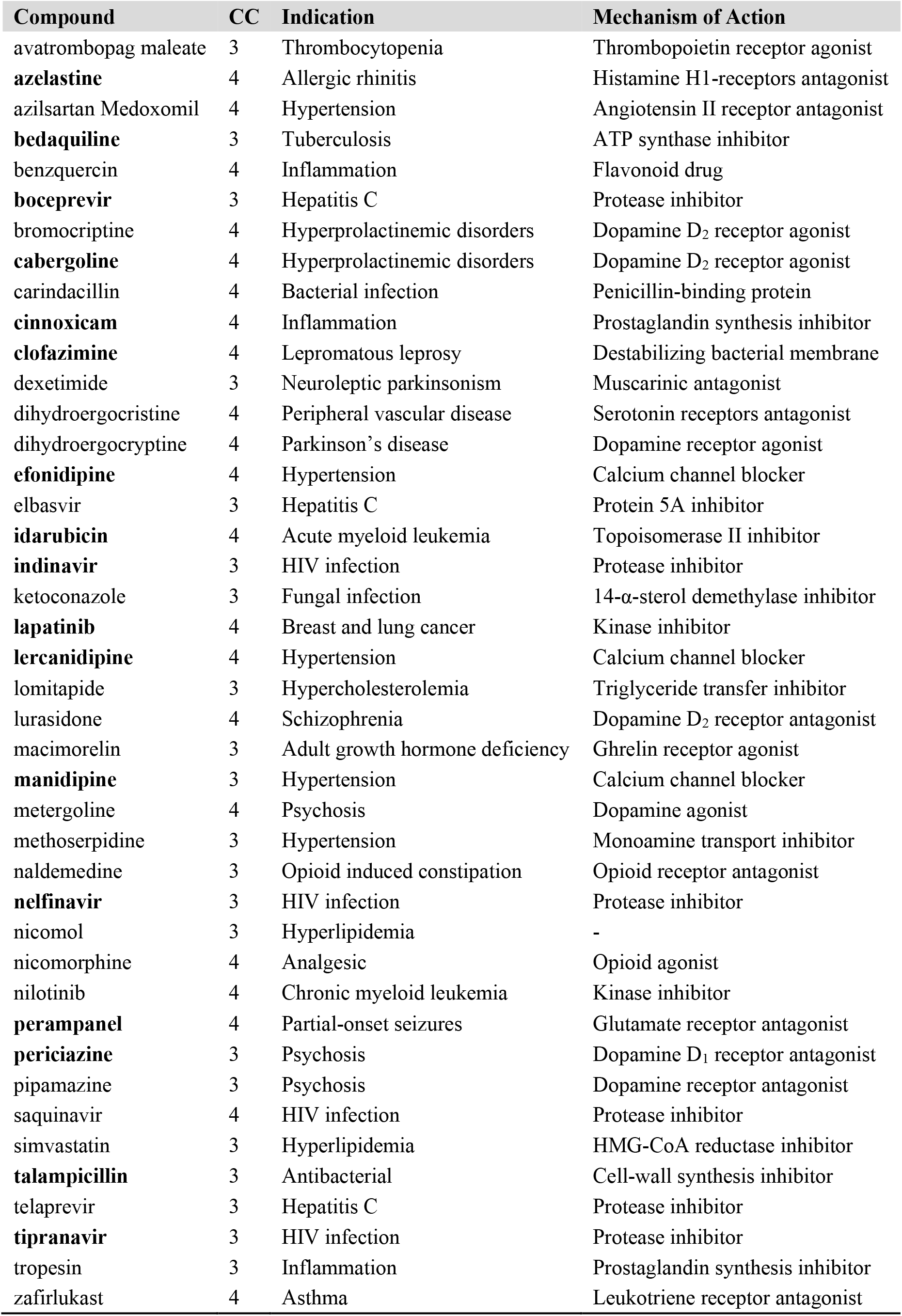
The Consensus Count (CC), Indication and Mechanism of Action of the Top 42 Drugs Selected from Virtual Screening. Assayed Compounds are in Bold.

**Figure 2.**
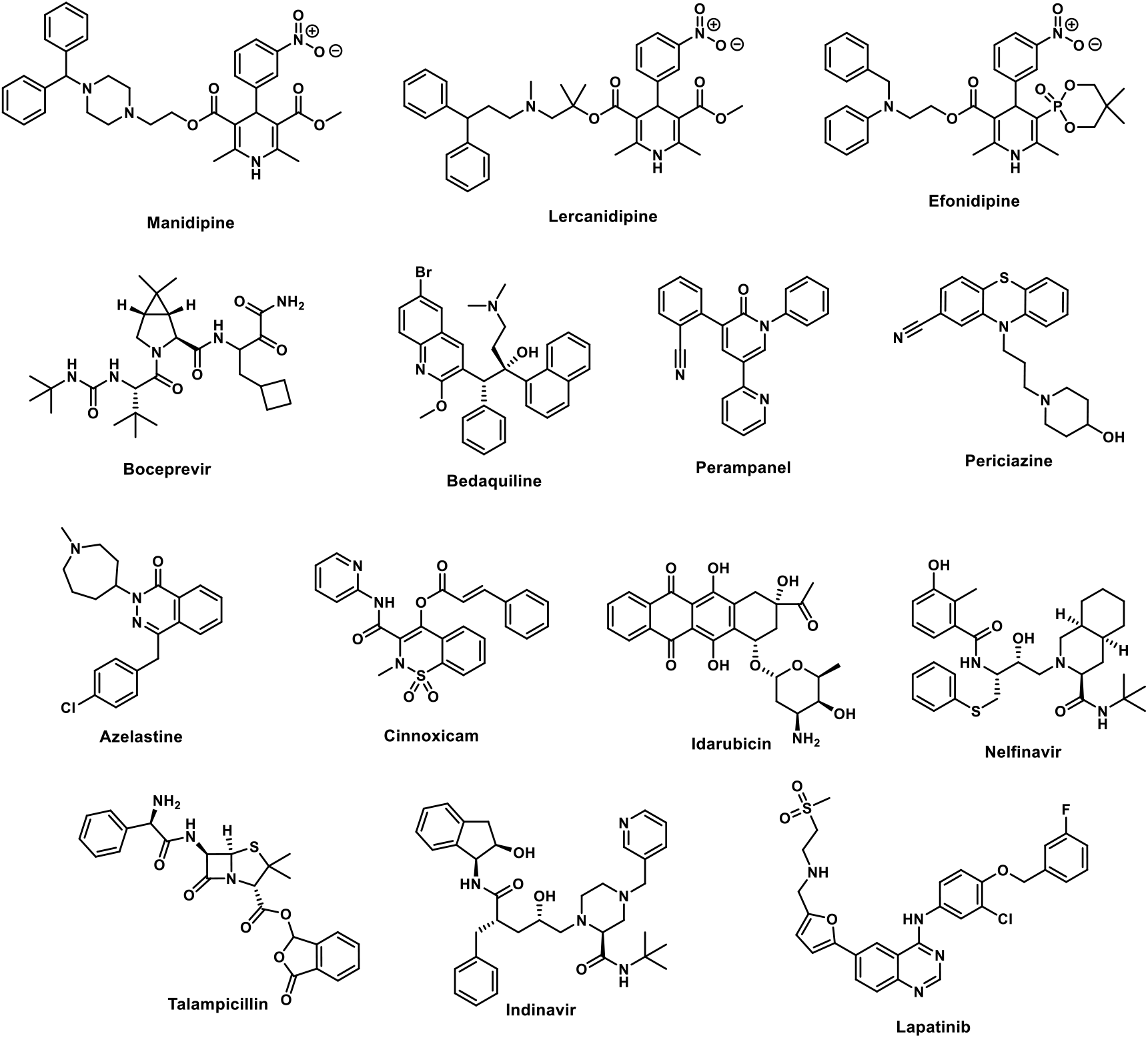
Selected high-scoring compounds from the consensus docking.

In almost all cases the predicted poses for the 42 compounds from the different docking programs agreed well. The poses from Glide were then subjected to extensive visual scrutiny to check for unsatisfied hydrogen-bonding sites, electrostatic mismatches, and unlikely conformation of the ligand. About half of the compounds were ruled out for further study due to the occurrence of such liabilities and the presence of multiple ester groups (e.g., methoserpidine and nicomol) or overall size and complexity (e.g., bromocriptine and benzquercin). A repeated motif was apparent with high-scoring ligands having a cloverleaf pattern with occupancy of the P1, P1’, and P2 pockets, as illustrated in Figure 3 for the complex of azelastine. Other common elements are an edge-to-face aryl-aryl interaction with His41 and placement of a positively-charged group in the P1 pocket in proximity to Glu166, e.g., the methylazepanium group of azelastine, the protonated trialkylamino group of bedaquiline, and protonated piperazine of periciazine. However, Glu166 forms a salt-bridge with the terminal ammonium group Ser1B (Figure 1). The electrostatic balance seems unclear in this region, so our final selections included a mix of neutral and positively-charged groups for the P1 site.

**Figure 3.**
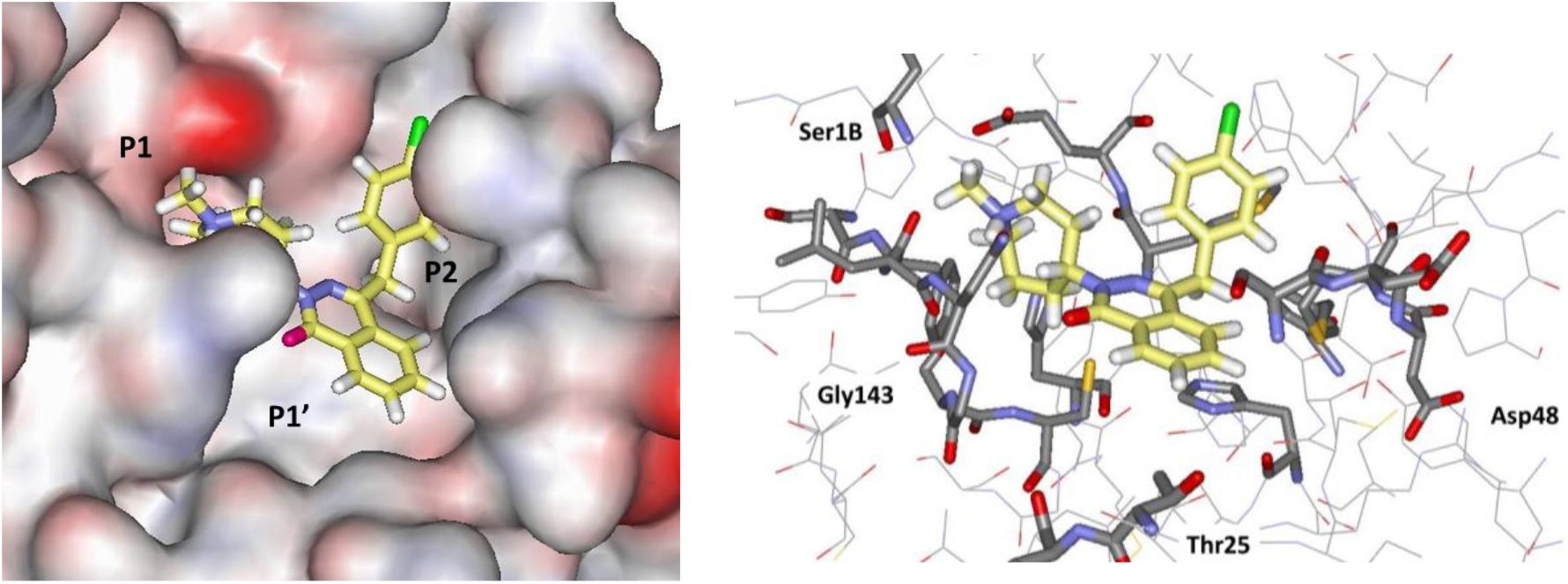
Glide docking pose for azelastine in space-filling (left) and stick (right) renderings. All illustrations are oriented with the P1 pocket to the left and P2 to the right, and all carbon atoms of ligands are in yellow.

The analysis of the high-scoring 42 compounds also considered structural variety and potential synthesis of analogs. In the end, we settled on 17 compounds, which are highlighted in Table 1, for purchase and assaying. Sixteen were commercially available, mostly from Sigma-Aldrich. The seventeenth, cinnoxicam, was not available, but it was readily prepared in a one-step synthesis from the commercially-available ester components. It may be noted that three calcium channel blockers, efonidipine, lercanidipine, and manidipine were purchased (Figure 2). This was not done owing to the characteristic dihydropyridine substructure, since this end of the molecule protrudes out of the P1’ site in the docked poses. It was for the variety in the left-sides of the molecules in Figure 2, which form the cloverleaf that binds in the P1, P1’, and P2 pockets, as illustrated in Figure 4 for manidipine. The steric fit in this region appears good, though the only potential hydrogen bond is between the nitro group and the catalytic Cys145.

**Figure 4.**
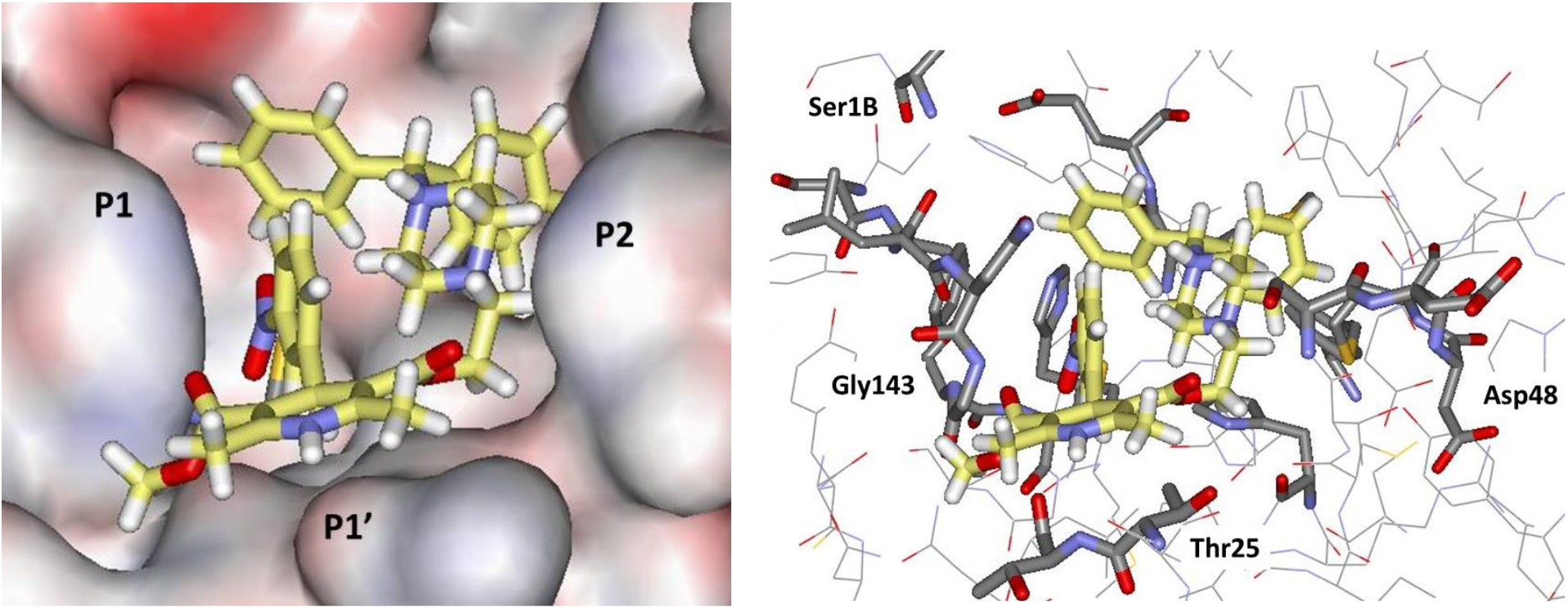
Glide docking pose for manidipine in space-filling (left) and stick (right) renderings.

### Protease Assay Results

The 17 known drugs were screened using the FRET-based assay monitoring the fluorescence generated from the cleavage of a peptide substrate harboring an Edans-Dabcyl pair by recombinant SARS-CoV-2 M^pro^. Remarkably, fourteen of the drugs at 100 μM decreased M^pro^ activity (100 nM), as shown in Figure 5 and Table 2. Five drugs decreased M^pro^ activity to below 40%. The top five hits from the kinetic assay were manidipine, boceprevir, efonidipine, lercanidipine, and bedaquiline. Dose-response curves were obtained to determine IC_50_ values, when possible, as shown in Figure 6 for the five most potent inhibitors, with the raw data as a function of time and concentration given in Figure S1.

**Figure 5.**
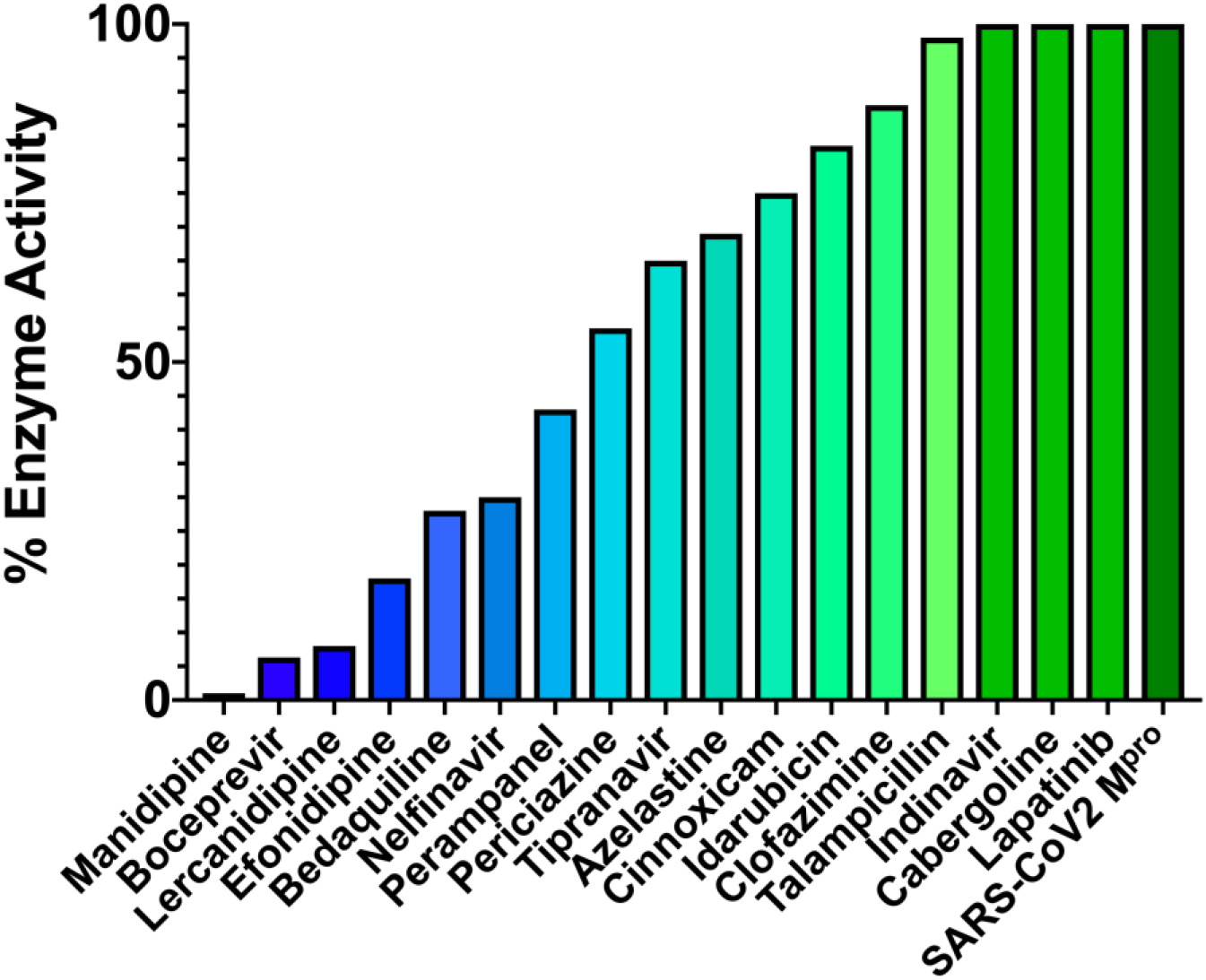
Ranking of the 17 compounds by percent residual enzyme activity monitored by cleavage product fluorescence following a one-hour incubation of 100 nM M^pro^ with 100 μM compound. Compounds are ranked from most (blue) to least (green) active.

**Table 2.** Measured Activities of the 17 Compounds Tested for Inhibition of M^pro^

| Compound | % Activity at 100 μM | IC<sub>50</sub> (μM) |
| --- | --- | --- |
| manidipine | 1 | 4.81 ± 1.87 |
| boceprevir | 6 | 5.40 ± 1.53 |
| lercanidipine | 8 | 16.2 ± 2.94 |
| efonidipine | 18 | 38.5 ± 0.41 |
| bedaquiline | 28 | 18.7 ± 4.20 |
| perampanel | 43 | 100-250<sup>a,b</sup> |
| periciazine | 55 | 250<sup>a</sup> |
| nelfinavir | 64 | 250-600<sup>a</sup> |
| tipranavir | 65 | >600<sup>a</sup> |
| azelastine | 69 | 20-100<sup>a</sup> |
| cinnoxicam | 75 | >600<sup>a</sup> |
| idarubicin | 82 | 250-600<sup>a</sup> |
| clofazimine | 88 | >600<sup>a</sup> |
| talampicillin | 90 | 250-600<sup>a</sup> |
| indinavir | 100 | NA |
| cabergoline | 100 | NA |
| lapatinib | 100 | NA |

<sup>a</sup> Estimate due to incomplete inhibition at 600 μM. <sup>b</sup> Fluorescence of compound interfered with assay.

**Figure 6.**
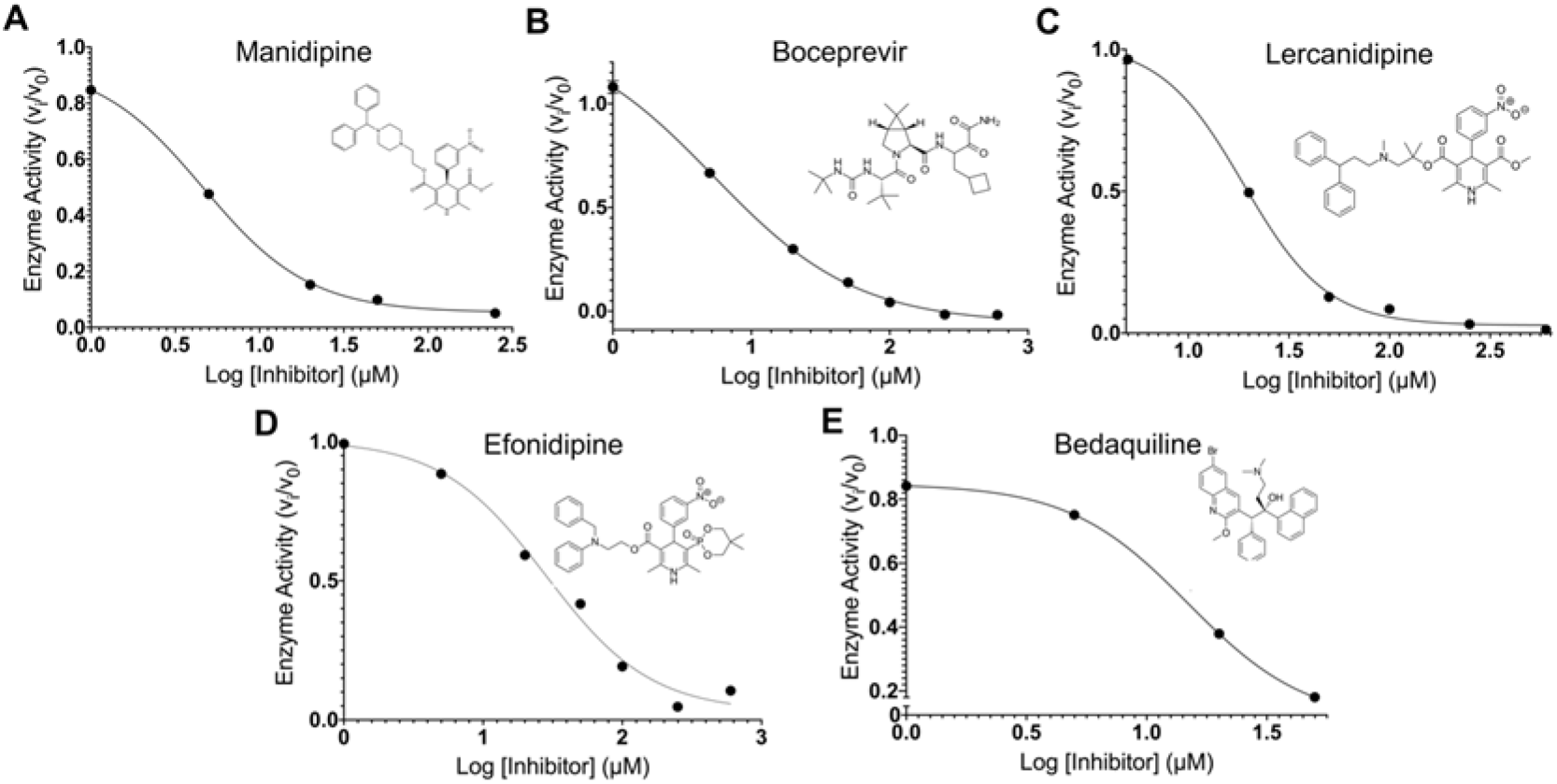
IC_50_ plots and values for the top five compounds active against SARS-CoV-2 M^pro^ from *in vitro* FRET-based assay. IC_50_ plots were generated from averaged kinetic data in triplicates for (A) manidipine, (B) boceprevir, (C) lercanidipine, (D) efonidipine, and (E) bedaquiline.

The calcium channel-blockers manidipine, lercanidipine, and efonidipine inhibit M^pro^ activity with IC_50_ values of 4.8 μM, 16.2 μM, and 38.5 μM, respectively. As suggested from Figure 4, the variation likely arises primarily from differences in binding of the left sides of the molecules (Figure 2) in the P1, P1’, and P2 pockets. It has previously been proposed that such compounds might be useful for treatment of SARS-CoV-2 infection for their role as calcium channel blockers, not as M^pro^ inhibitors.^44^ Boceprevir, a hepatitis C virus protease inhibitor, inhibits M^pro^ with an IC_50_ of 5.4 μM; its IC_50_ has been previously reported as 4.13 μM.^7^ Bedaquiline, approved for the treatment of multi-drug-resistant tuberculosis, inhibits M^pro^ with an IC_50_ of 18.7 μM. The IC_50_ of nelfinavir, an HIV protease inhibitor, was estimated to be between 250 and 600 μM. Vatansever et al. have previously reported an IC_50_ for nelfinavir of 234 μM.^45^ Perampanel appears to be the sixth most active compound at 100 μM, though its IC_50_ could not be calculated reliably, as its intrinsic fluorescence interfered with the fluorescence measurements.

The computed structures for the complexes of boceprevir and bedaquiline are illustrated in Figure 7. For boceprevir, the dimethylcyclopropyl subunit is predicted to sit in P1, the sidechain with the cyclobutyl and terminal ketoamide groups is in P1’, the t-butyl group is in P2, and there are hydrogen bonds with the NH of Gly143 and carbonyl oxygen of Thr26. For bedaquiline, the three pockets are occupied by the ammonium containing sidechain, the naphthyl group, and the phenyl group, respectively, while the quinoline fragment extends towards the solvent, and there are no clear protein-ligand hydrogen bonds. The activity of this compound does suggest that positively-charged groups may be acceptable in the P1 site.

**Figure 7.**
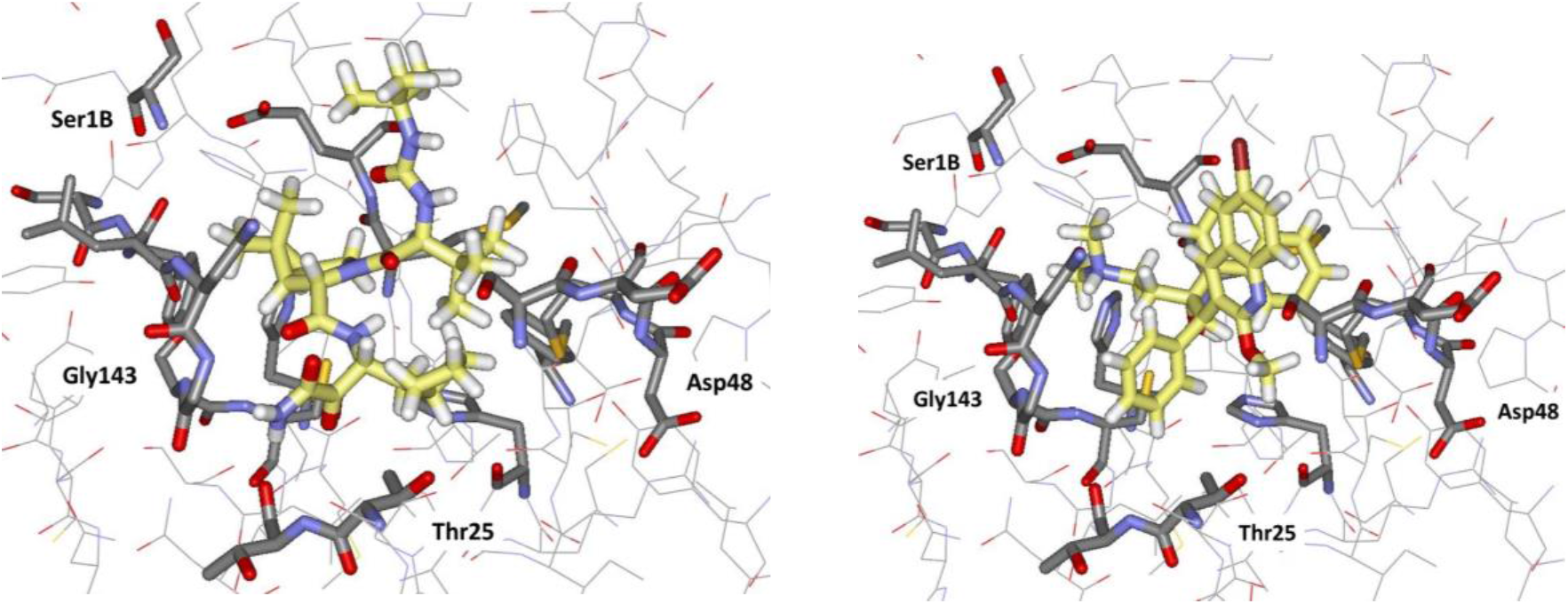
Renderings of Glide docking poses for (left) boceprevir, and (right) bedaquiline.

### MD Analyses for the M^pro^-ligand Complexes

Before the assaying was carried out, the 70-ns MD simulations were run for complexes of 14 of the promising compounds starting from the Glide poses. The idea was to obtain insight on which compounds gave more stable complexes and were, therefore expected to be more active inhibitors. In addition to visualization of the evolving structures, the all-atom RMSD of each ligand was computed over the course of the simulation time with and without least-square (LS) fitting of the ligand’s atoms onto the initial orientation of the complex (Figure 8). The LS-fit RMSD monitors only the ligand’s conformational changes, whereas the no-fit RMSD also reflects rotational and translational movements. The LS-fit RMSD converged relatively quickly to 2-3 Å for all ligands, except for carindacillin, which converged to 4 Å. However, as expected, no-fit RMSD values are larger than the LS-fit RMSD values for all ligands, demonstrating the contribution of rotational and translational movements. The no-fit RMSD value converged for bedaquiline, idarubicin, indinavir, and perampanel after about 10 ns, while it converged for efonidpine after 60 ns. The no-fit RMSD values steadily fluctuated about an average value of 4 Å for lapatinib and periciazine. In general, most ligands showed some displacement from their initial position, whereas they remained close to their initial conformation. Clear correlation of the results with the measured activities is not obvious, perhaps because 70 ns is too short a timeframe. For example, the RMSDs for bedaquiline and perampanel converge well, while they are more erratic for the more active boceprevir and efonidipine. More sophisticated MD procedures for gauging stability are known such as metadynamics, steered-MD, and random-accelerated MD,^46–49^ which would be interesting to apply retrospectively to the present experimental results.

**Figure 8.**
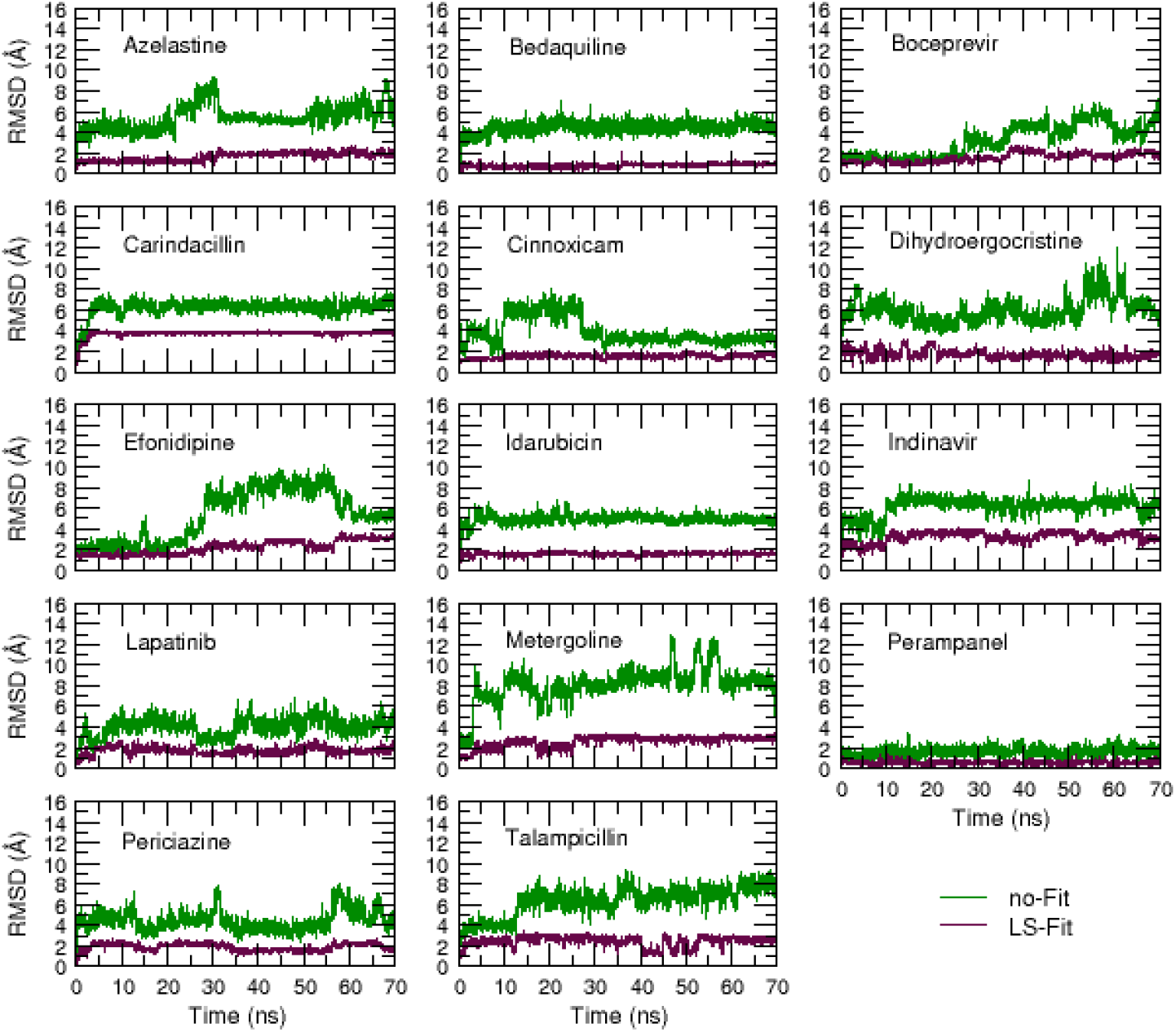
RMSD in Å of all ligand atoms with and without Least-Square fitting to the original complex structure during the course of 70-ns MD simulations.

## CONCLUSIONS

The present virtual screening study was highly successful in identifying 14 known drugs as showing inhibitory effect on the main protease of SARS-CoV-2. The consensus scoring approach using three docking programs and four protocols was effective in narrowing down ca. 2000 candidate drugs to 42 of high interest. The final 17 compounds that were selected for assay did reflect additional human visualization and analyses, though assaying of all 42 top compounds would not be burdensome. Five compounds were identified with IC_50_ values below 40 μM with manidipine, boceprevir, lercanidipine, and bedaquiline having values of 4.8, 5.4, 16.2, and 18.7 μM. Further study of these compounds in the context of COVID-19 therapy is warranted, while all of the active compounds reported here may provide a foundation for lead optimization to deliver valuable chemotherapeutics to combat the pandemic.

## Supporting information

Docking Scores

## ASSOCIATED CONTENT

### Supporting Information

The Supporting Information is available free of charge on the ACS Publications website. An Excel file with the names and docking scores for the full drug library, a Figure with kinetic data for the assays of the five most active compounds, and a Figure comparing docking scores.

## AUTHOR INFORMATION

### Notes

The authors have no competing interests.

## ACKNOWLEDGMENTS

Gratitude is expressed for support to the U. S. National Institutes of Health (GM32136) and to the Yale University School of Medicine for a CoReCT Pilot Grant. The M^pro^ plasmid was kindly provided by the Hilgenfeld lab.^5^

