## Supplementary Figures for "Identification of 14 Known Drugs as Inhibitors of the Main Protease of SARS-CoV-2"

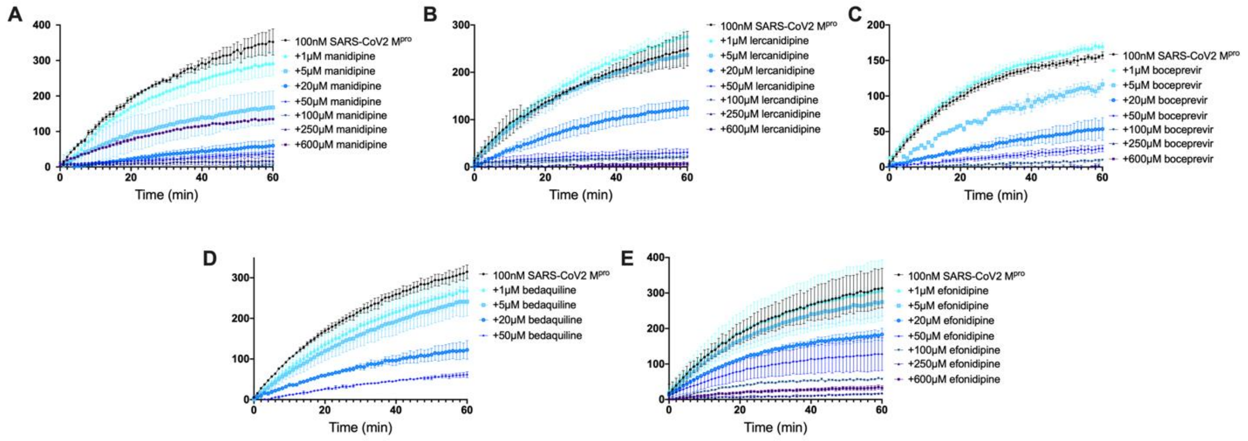


**Figure S1.** Kinetic data measuring activity of SARS-CoV2 $M^{pro}$ in the presence of A) manidipine, B) lercanidipine, C) boceprevir, D) bedaquiline, and E) efonidipine. All measurements were performed in triplicate, averaged, and plotted with standard deviation. Kinetic data for inhibition of $M^{pro}$ by bedaquiline was collected at concentrations up to 50 μM due to solubility issues.

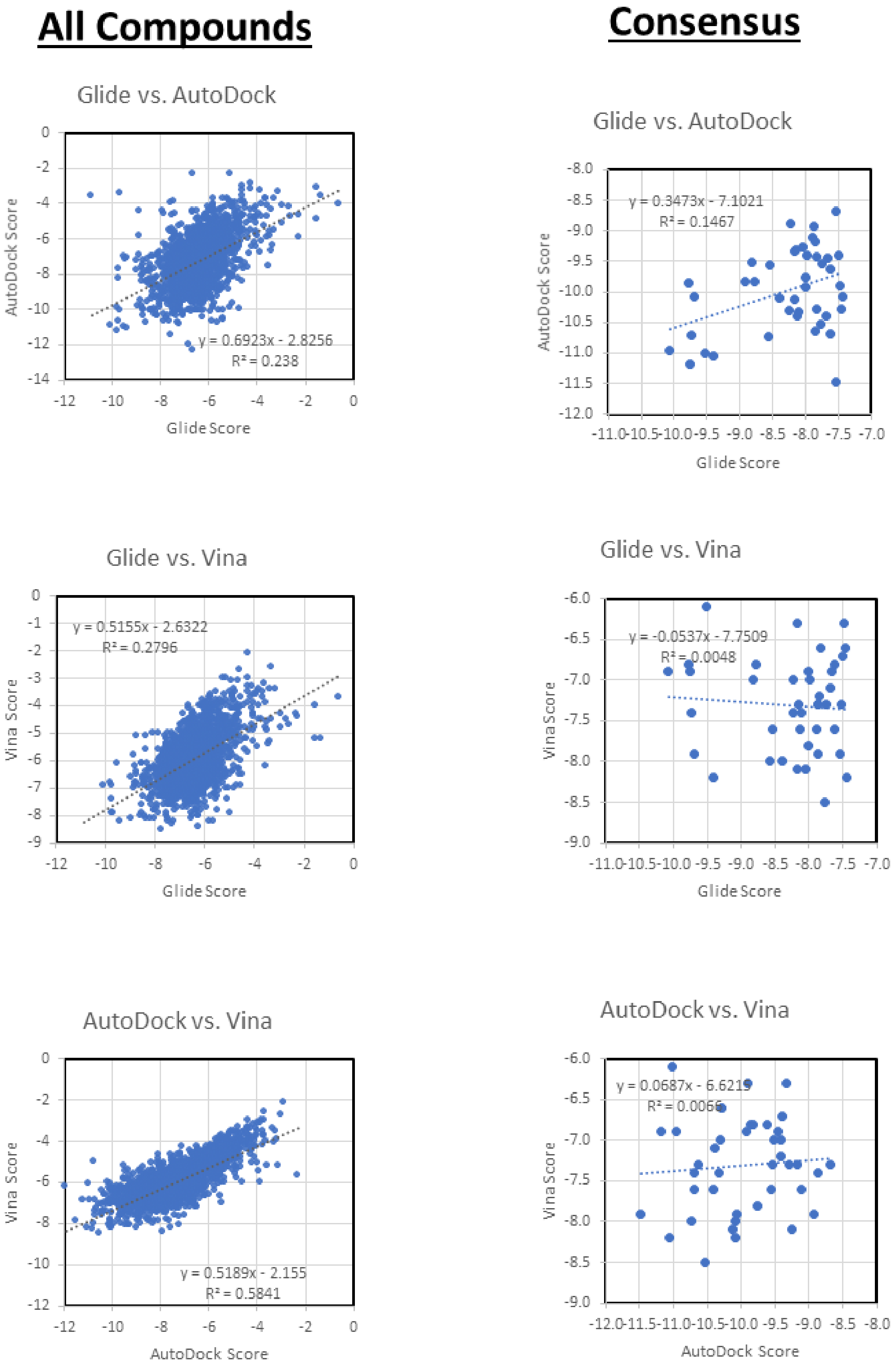


**Figure S2.** Correlation of docking scores for all drugs (left) and for the 42 consensus compounds (right).
